# HyperSketch: *de Bruijn* graph sketching for genomic similarity estimation with Hyperdimensional Computing

**DOI:** 10.64898/2026.09.01.748726

**Authors:** Fabio Cumbo, Kabir Dhillon, M. Hassan Najafi, Sercan Aygun, Daniel Blankenberg

## Abstract

**Background:** The exponential growth of genomic databases necessitates alignment-free methods for comparing genomes. While MinHash-based tools have revolutionized this field by efficiently estimating the Average Nucleotide Identity based on k-mer sets, they inherently discard structural genomic information.

**Methods:** We introduce HyperSketch, a novel sketching tool that encodes the *de Bruijn* graph structure of a genome into a fixed-size, topology-aware *vector* using Hyperdimensional Computing (HDC). Unlike set-based sketches, HyperSketch encodes the transitions between adjacent k-mers into a superposition of orthogonal hypervectors. To formalize parameter selection, we also propose an analytical framework proving that graph-based sketches fundamentally require a smaller k-mer size than set-based models due to their expanded k+1 biological footprint.

**Results:** We benchmarked HyperSketch against Mash and HyperGen using a dataset of ∼26 thousand viral reference genomes from NCBI GenBank. Under optimal parameters, we demonstrate a strong linear correlation (>99%) between HyperSketch’s graph-based similarity and standard MinHash distance estimates. Crucially, we show that the mathematical formulation of HyperSketch introduces a distance scaling effect that expands the dynamic range of estimates for closely related strains, providing a higher-resolution metric for sub-lineage clustering than purely compositional estimators.

**Conclusions:** HyperSketch provides a computationally efficient, structure-aware alternative to traditional sketching. By natively encoding genomic syntax, it offers a new dimension of genomic comparison that excels at both high-resolution strain differentiation and deep evolutionary scaling, complementing existing nucleotide identity metrics without requiring sequence alignment.

## INTRODUCTION

The last two decades have witnessed a sharp rise in genomic data, driven by the rapid advancement and decreasing cost of high-throughput sequencing technologies [1]. Public repositories such as the National Center for Biotechnology Information (NCBI) and the European Nucleotide Archive (ENA) are now accumulating petabytes of genomic sequences at an exponential rate [2]. While this wealth of data offers extraordinary opportunities for comparative genomics, evolutionary biology, and large-scale pathogen surveillance, it also introduces a severe computational bottleneck [3].

Traditional sequence comparison methodologies rely on alignment algorithms (such as BLAST [4] or MUMmer [5]) that utilize dynamic programming or exact substring matching. Although these alignment-based methods represent the gold standard for base-level accuracy and homology detection, their quadratic or super-linear time and space complexities render them computationally intractable for massive, dataset-scale comparisons [6]. As the sheer volume of sequenced genomes continues to outpace Moore’s Law, relying on alignment for preliminary genomic distance estimation or all-*vs*-all database clustering is no longer feasible. Consequently, there is an urgent need for highly scalable computational methodologies capable of rapidly estimating genomic similarity without the computational overhead of exact sequence alignment.

To circumvent the computational burdens of exact alignment, the bioinformatics community has increasingly turned to alignment-free methodologies [7–9]. These approaches predominantly rely on decomposing sequences into sets of overlapping substrings of length k, known as k-mers. By evaluating the overlap of k-mer profiles, researchers can estimate the evolutionary distance between organisms. However, as datasets grow, storing and intersecting exhaustive k-mer sets for thousands of genomes still requires prohibitive amounts of memory [10].

To resolve this, locality-sensitive hashing and dimensionality reduction techniques, broadly referred to as sketching, have been widely adopted. The most transformative of these algorithms is MinHash [11,12], which was successfully popularized in genomics by tools such as Mash [13]. Instead of storing the entire k-mer vocabulary, Mash hashes the constituent k-mers and retains only a fixed-size subset of the smallest hash values. By merely comparing these highly compressed sketches, the algorithm rapidly approximates the Jaccard similarity index [14] of the original genomes. This Jaccard index is then mathematically converted into an estimation of the Average Nucleotide Identity (ANI). By drastically reducing the genome size, MinHash-based estimators have substantially reduced sequence comparison times from CPU-hours to milliseconds, enabling the routine clustering and querying of massive global databases.

Despite the immense computational advantages of MinHash and other set-based sketching algorithms, they share a fundamental algorithmic limitation: *they model genomes as unstructured bags of words* [13,15–17]. By reducing a biological sequence to an isolated set of k-mers, these methods inherently sever the contextual links between adjacent elements. Consequently, set-based estimators are entirely blind to genomic syntax and higher-order structural organization. This limitation becomes particularly problematic when analyzing structural variants, such as large-scale inversions, horizontal gene transfers, or recombination events, phenomena that are heavily prevalent in viral evolution. For instance, if a viral genome undergoes a perfect inversion of a major gene segment, its overall k-mer composition remains virtually identical, save for a few disrupted k-mers at the breakpoints. A standard MinHash algorithm will evaluate the original and inverted genomes and report a distance approaching zero, falsely implying near-perfect homology. To accurately resolve complex evolutionary trajectories and differentiate between compositionally similar but structurally rearranged strains, comparative tools must move beyond simple set intersections and incorporate the topological order of the genome.

To capture this essential structural information without reverting to the computational expense of sequence alignment, we turn to Hyperdimensional Computing (HDC) [18,19], also known as Vector Symbolic Architectures (VSA), which has recently gained significant attention across the biomedical sciences as a highly-efficient, noise-tolerant computing paradigm [20,21]. Inspired by the distributed representation of information in the human brain, HDC computes using massive, high-dimensional *pseudo*-random vectors, typically containing thousands of dimensions (e.g., D=10,000; denoted as *hypervector*). In this high-dimensional space, the geometry dictates that any two randomly generated vectors are nearly orthogonal to one another. The true power of HDC lies in its well-defined algebraic operations, specifically *binding* (element-wise multiplication in bipolar data +1, -1; or XOR in binary data 1, 0) and *bundling* (element-wise addition or superposition). By binding vectors representing distinct elements, HDC can cleanly encode relationships, sequences, and graph edges. By bundling these bound vectors, it can superimpose thousands of complex relationships into a single, fixed-size holographic vector. Furthermore, because information is distributed holistically across all dimensions rather than localized to specific bits, HDC representations are inherently robust to extreme quantization, noise, and hash collisions, making it an ideal mathematical framework for compressing massive relational datasets.

The application of HDC to genomic sketching has been recently pioneered by tools such as HyperGen [22], which successfully demonstrated that hypervectors can approximate Jaccard similarities with extreme memory efficiency. However, HyperGen fundamentally operates under the same bag-of-words paradigm as Mash. It encodes isolated k-mers into hypervectors without preserving their genomic syntax. Consequently, while it leverages the computational benefits of HDC, it inherits the exact same structural blindness as traditional MinHash estimators.

In this study, we introduce HyperSketch, a novel computational framework that bridges the gap between high-speed genomic sketching and structural awareness. Instead of discarding positional context to create a set-based bag-of-words, HyperSketch constructs an implicit *de Bruijn* sequence graph and directly encodes its topological edges, i.e., the transitions between adjacent kmers [22–24]. Utilizing HDC encoding and operations, these structural transitions are deterministically mapped, bound, and bundled into a fixed-size, highly compressed memory matrix. This approach allows HyperSketch to achieve the extreme dimensionality reduction and computational efficiency characteristic of traditional MinHash estimators, while simultaneously preserving a robust, holographic representation of the genome’s structural syntax. Consequently, HyperSketch intrinsically penalizes complex structural rearrangements, providing a rapid, topology-aware metric for evolutionary distance estimation that complements existing compositional analyses.

## MATERIALS AND METHODS

The HyperSketch framework is designed to transform raw genomic sequences into highly compressed, structure-aware hyperdimensional sketches. The code algorithm, implemented in C++, operates in four sequential stages: (i) implicit *de Bruijn* graph extraction from raw FASTA sequences, (ii) hyperdimensional encoding of topological edges into a dense Memory Bank, (iii) extreme sketch compression via bit-packing and algorithmic bias mitigation, and (iv) structural distance estimation utilizing geometric similarity metrics. Furthermore, to adapt this architecture specifically for viral comparative genomics, we outline both the theoretical models and empirical frameworks used to optimize the algorithm’s sequence footprint.

### Sequence Parsing and *de Bruijn* Graph Extraction

The HyperSketch encoding pipeline begins by reading a genomic sequence and implicitly constructing its underlying *de Bruijn* graph. The nucleotide sequence is decomposed into a rolling window of overlapping k-mers. To ensure the sketch is independent of the sequencing strand (*forward* or *reverse*), each extracted sequence is canonicalized.

Unlike set-based estimators that store the canonicalized k-mer as an isolated node, HyperSketch links adjacent canonical k-mers to define a topological edge. To maintain strict strand-independence and allow HyperSketch to perfectly match reverse-complemented contigs, the graph is treated as undirected. The nodes constituting the edge are lexicographically sorted prior to encoding. This extraction formally models the genome as a set of structural transitions. If a non-ACGT character (e.g., an N representing an ambiguous base or a scaffold gap) is encountered, the rolling window is instantly reset. This prevents the algorithm from falsely bridging distant genomic regions, strictly preserving the true biological topology of the contigs.

### Hyperdimensional Encoding and the Memory Bank

To compress the extracted graph topology into a fixed-size representation, HyperSketch maps the sequences of undirected edges into a high-dimensional associative memory array, referred to as the Memory Bank (M). The mathematical foundation of HDC relies on the geometry of high-dimensional spaces (e.g., D=10,000). In such spaces, any two randomly generated bipolar vectors are nearly orthogonal (i.e., the cosine similarity approaches zero). This pseudo-orthogonality allows HDC to aggregate multiple vectors into a single composite representation without entirely destroying the individual signals.

To manipulate these representations, HDC utilizes the foundational MAP (Multiply-Add-Permute) operational model. The MAP framework defines three core algebraic HDC operations: Multiplication (binding) to associate distinct concepts, Addition (bundling) to superimpose multiple vectors into an aggregated macroscopic set, and Permutation to encode strict sequence or order.

HyperSketch utilizes a matrix M of dimensions *M_rows_* × *D*, where *M_rows_* represents the number of available hash buckets (e.g., *M_rows_* = 4, 000) and *D* represents the hypervector dimensionality. Instead of utilizing traditional multiplicative HDC binding to link nodes, HyperSketch adapts the MAP framework by employing a highly efficient spatial *key*-*value* binding paradigm. For each extracted undirected edge (source → target), the algorithm performs a sequential, three-step encoding process:

1. <u>Deterministic projection:</u> the target node is mapped to a unique pseudo-orthogonal hypervector. To ensure reproducibility across independent executions, meaning identical k-mers will always generate identical hypervectors across different executions, the hypervector is generated deterministically. The string representation of the target node is hashed into a 64-bit integer. This integer directly seeds a Weyl-sequence pseudo-random number generator (PRNG) [25], which rapidly populates the D-dimensional vector with uniformly distributed bipolar numbers.
2. <u>Spatial Addressing (binding):</u> rather than generating a second hypervector for the source node, the source sequence acts as the addressing key for the Memory Bank [26]. A rapid hashing function computes the row index for the source node. Because *M_rows_* is exactly bounded and orders of magnitude smaller than the total combinatorial space of all possible k-mers, this spatial addressing enforces extreme dimensionality reduction.
3. <u>Holographic Superposition (bundling):</u> Once the destination row is identified by the source hash, the generated target hypervector is element-wise added (bundled) into the selected row. During the encoding phase, the Memory Bank is maintained as an array of 32-bit integers to accurately accumulate the superimposed values without early saturation.

Because the number of unique source nodes in a viral genome vastly exceeds *M_rows_*, hash collisions are inevitable. Multiple distinct source nodes will frequently map to the exact same row in the Memory Bank. In HDC, the row becomes a holographic superposition of all topological transitions originating from that bucket.

When two distinct starting sequences end up assigned to the exact same location in the memory bank, their corresponding target sequences are simply merged together into a single combined representation. If this sketch is later compared against another genome that only shares one of those original connections, the unrelated overlapping data simply acts as background noise. Because the target hypervectors are sampled independently from a uniform bipolar distribution, their superposition behaves as zero-mean interference. When the similarity between two sketches is later calculated via dot product, the matching sequence topologies constructively interfere to produce a strong geometric signal. Simultaneously, the uncorrelated mismatched vectors exhibit destructive interference, with their inner products statistically aggregating to zero. Consequently, HyperSketch can compress the vast, highly sparse network of sequence transitions into a dense, fixed-size fingerprint without losing the underlying structural data.

Crucially, this holographic superposition ensures that structural features are not stored as isolated, separate facts. Instead, they are structurally integrated into a single, indivisible composite representation, meaning any variation in the underlying *de Bruijn* graph topology geometrically alters the entire hypervector.

### Sketch Compression and Binarization

During the encoding phase, the Memory Bank M utilizes standard integers to accurately accumulate the superimposed vectors. To minimize the storage footprint of the final sketch, the matrix is aggressively quantized into a binary format. Each element in the matrix is reduced to a single bit based on its sign.

Statistically, due to the symmetrical distribution of bundled pseudo-random vectors, sums that equal exactly zero occur frequently. Assigning these zero-sums uniformly to a single binary class introduces a systematic zero-bias that artificially inflates the similarity between unrelated sketches. To eliminate this algorithmic artifact, HyperSketch applies a deterministic cryptographic tie-breaker. For each active memory bucket, a unique 64-bit seed is generated by hashing the absolute values and positional indices of its non-zero dimensions using an FNV-1a offset basis and SplitMix64 bit-mixing. This deterministic seed initializes a stateless SplitMix64 pseudo-random number generator (PRNG) utilizing a Weyl sequence, which assigns consistent, pseudo-random bits to the zero-sum dimensions. This guarantees that genomes with identical structural sub-graphs receive identical tie-breakers, while divergent graphs receive uncorrelated noise. This strictly maintains a 50/50 probability distribution for zero-sums. Finally, the binarized matrix is bit-packed (storing 8 dimensions per byte), achieving an exact 32:1 compression ratio over the integer matrix.

The complete step-by-step process for generating a compressed sketch from a raw genome sequence is detailed in Algorithm 1.

**Algorithm 1**: Pseudocode detailing the complete execution path for transforming a raw genomic sequence into a compressed, structure-aware hyperdimensional sketch. The pipeline encompasses implicit *de Bruijn* graph extraction, deterministic HDC projection, vector bundling, and zero-bias mitigated binarization.

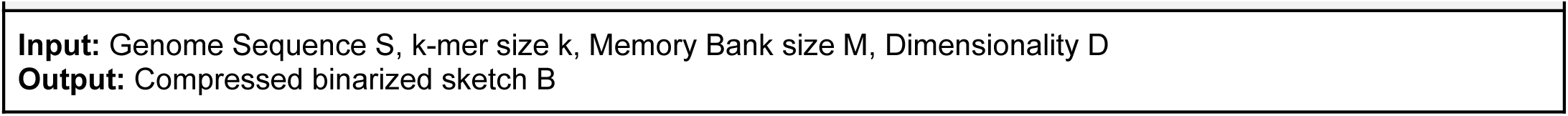
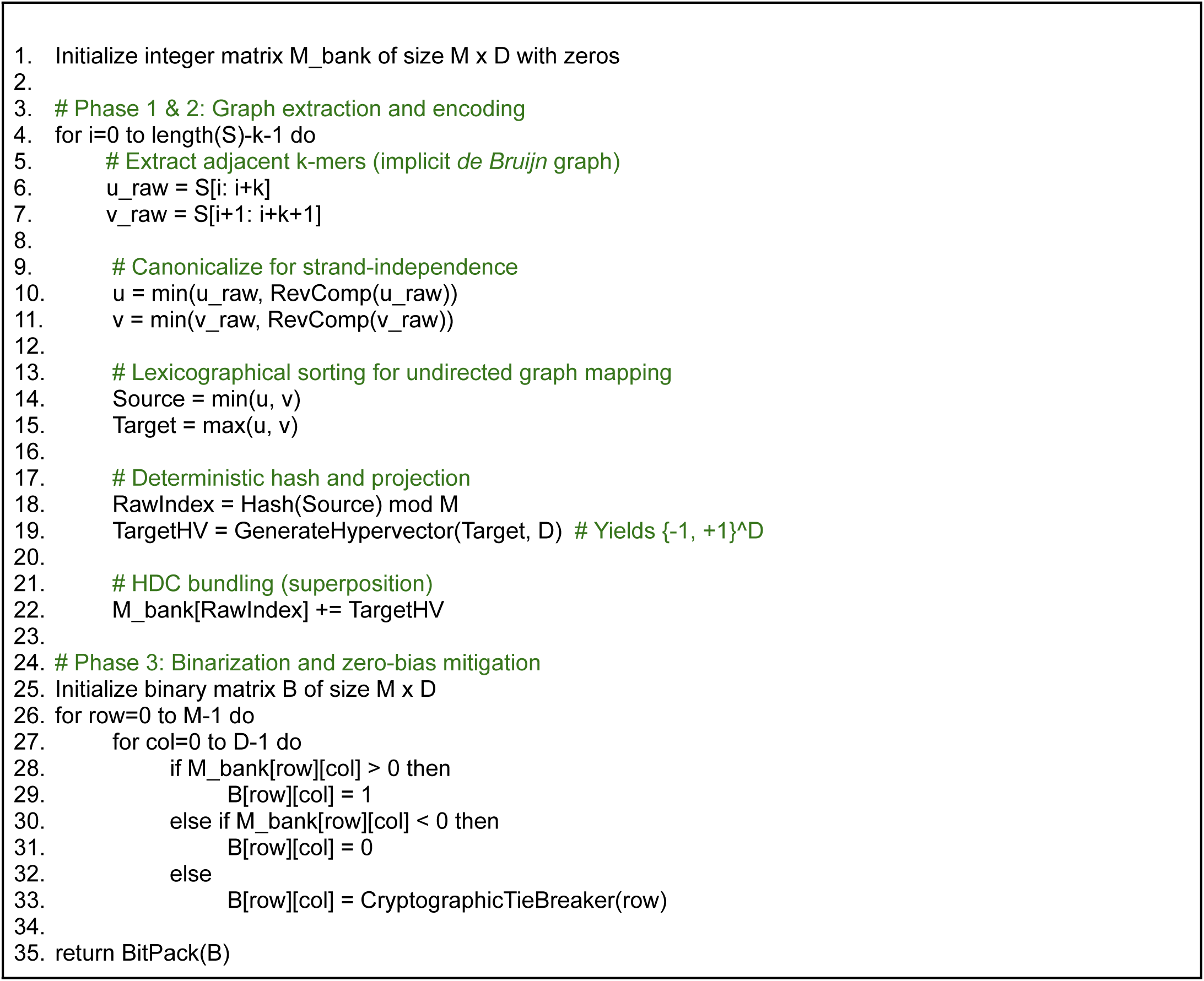

### Distance Calculation and Edge-Footprint Adjustment

To compare two binary sketches A and B, we compute the Binarized Cosine Similarity. To mitigate the sparsity of the memory bank M, the denominator utilizes the geometric magnitude of active buckets rather than the Jaccard union:

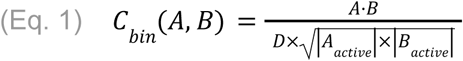

Directly utilizing this binarized similarity is insufficient, as aggressively quantizing massive integers into single bits produces a geometric distortion that warps the perceived relationship between sketches. To correct the geometric distortion introduced by aggressive quantization, we apply Grothendieck’s identity [27] to recover the continuous cosine similarity *C* from the binarized cosine similarity *C_bin_*, mathematically defined as:

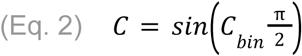

This step seamlessly estimates what the true, continuous similarity would have been had the Memory Bank remained entirely uncompressed.

Once the true continuous similarity is recovered, we apply the Otsuka-Ochiai transformation [28]. Because HyperSketch measures the geometric angle between vectors, while traditional mutation models rely on the fractional overlap of discrete sets, this transformation acts as a crucial analytical bridge. It explicitly converts the vector-based cosine similarity into an equivalent Jaccard index (J), allowing the continuous hyperdimensional output to be treated exactly like a traditional k-mer overlap:

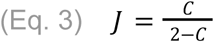

Finally, the estimated Jaccard index *J* is transformed into a genomic distance. Because HyperSketch operates on structurally embedded *de Bruijn* edges with a biological footprint of k+1 nucleotides, the standard Mash transformation is mathematically adapted to:

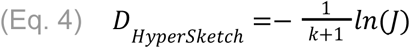

This formulation strictly reflects the topological decay rate of these expanded features under mutation, bypassing the standard set-based assumptions.

### Theoretical and Empirical Parameter Optimization

The selection of the k-mer size in HyperSketch dictates the resolution of the underlying *de Bruijn* sequence graph. Because HyperSketch encodes topological edges (*k_i_* → *k_i_*_+1_) rather than isolated nodes, the effective biological footprint of a single encoded feature is k+1 nucleotides. To determine the optimal k-mer size for graph-based sketching, we developed a comprehensive theoretical and empirical modeling framework that balances two opposing topological boundaries.

### Theoretical Topological Viability

The optimization of a graph sketch is governed by a fundamental trade-off between edge uniqueness and edge survival. First, the combinatorial space of all possible edges (4) must be sufficiently larger than the genome length (L) to prevent unrelated genomes from sharing graph edges purely by random chance (homoplasy). We model the probability of Edge Uniqueness as:

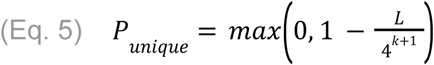

Second, as k increases, the sequence graph becomes increasingly fragile. A single nucleotide substitution within the k+1 footprint will completely sever the topological transition between the two nodes. Given an evolutionary mutation rate *p*, the probability of Edge Survival is modeled as:

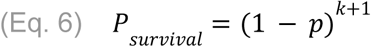

We formally define Topological Viability (V) as the product of these two opposing probabilities:

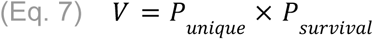

This theoretical model demonstrates that because a graph edge requires k+1 intact bases to survive, it decays exponentially faster than an isolated node of length k. Consequently, a graph-based sketch fundamentally requires a smaller k-mer size than standard set-based (bag-of-words) models to maintain the same optimal signal-to-noise ratio.

#### Empirical Validation via Sequence Set Intersections

While theoretical viability models provide a foundational mathematical boundary, they rely on the heuristic assumption of a uniform nucleotide distribution. True viral genomes exhibit distinct GC-biases, repetitive motifs, and evolutionary constraints, causing random collision probabilities to derive from uniform models.

To account for these biological realities, we complemented our theoretical model with an empirical parameter sweep analysis. We took inspiration from the original Kitsune framework [29], which was developed to determine optimal k-mer lengths for alignment-free phylogenomics by estimating sequence uniqueness. To adapt this concept for exact structural topologies, we implemented a custom, highly optimized C++ evaluator that performs exact lock-free set intersections across our entire viral dataset. Rather than evaluating isolated k-mers, this engine strictly extracts and evaluates (k+1)-mers to mirror HyperSketch’s edge footprint. By empirically measuring both the true absolute sequence uniqueness and the number of average common features across varying k-mer sizes, this framework allowed us to confidently identify the parameter that maximizes the surviving homologous signal in our sequence graphs.

## RESULTS

In this section, we present the evaluation of HyperSketch on a comprehensive dataset of approximately 26,285 viral genomes from NCBI GenBank [30] organized into 17 phyla, 38 classes, 63 orders, 161 families, 1,014 genera, and 8,169 species. We first demonstrate, both theoretically and empirically, that sketching structural graph topologies fundamentally requires a smaller k-mer footprint than standard bag-of-words estimators to maximize the surviving biological signal. Utilizing these optimized parameters, we then benchmark HyperSketch’s genomic distance estimations against Mash, revealing a strong linear correlation across standard evolutionary relationships. Finally, we highlight HyperSketch’s unique capability to detect structural variations, successfully differentiating between compositionally identical but structurally rearranged genomes, a critical dimension of evolutionary distance that purely set-based estimators fail to capture.

### The Node vs. Edge Footprint Trade-off

To evaluate our theoretical models under biologically representative conditions, we established a baseline genome length (L) of 30 thousand base pairs. This length was selected as a representative standard, as it reflects the approximate size of several heavily sequenced viral families that dominate our set of 26 thousand viral genomes, like *Coronaviridae* (∼30kb) and *Adenoviridae* (∼35kb). While true viral genome sizes vary by orders of magnitude, establishing L=30,000 provides a realistic, intermediate combinatorial baseline for modeling the probability of random sequence collisions (homoplasy) in typical epidemiological datasets.

Alongside genome length, modeling the probability of topological survival requires a biologically appropriate range of evolutionary mutation rate [31]. We evaluated the models across a spectrum of divergence thresholds: 2% (representing closely related intra-species variants), 5% (a standard species-delimitation boundary corresponding to ∼95% Average Nucleotide Identity), and 10% (capturing deeper evolutionary divergence between distinct viral lineages).

Using these parameterized standards, theoretical modeling revealed a fundamental mathematical divergence between bag-of-words estimators and *de Bruijn* graph sketches. We evaluated the Topological Viability score, a metric balancing the need for combinatorial uniqueness against the probability of signal degradation under evolutionary pressure, across varying k-mer footprints.

The optimization curves in Figure 1 expose why traditional MinHash parameter defaults are suboptimal for structural graph encoding. Because an implicit *de Bruijn* edge encompasses k+1 intact bases to survive a point mutation (compared to only k bases for an isolated MinHash node), sequence graph topologies shatter exponentially faster under equivalent mutation rates.

**Figure 1:**
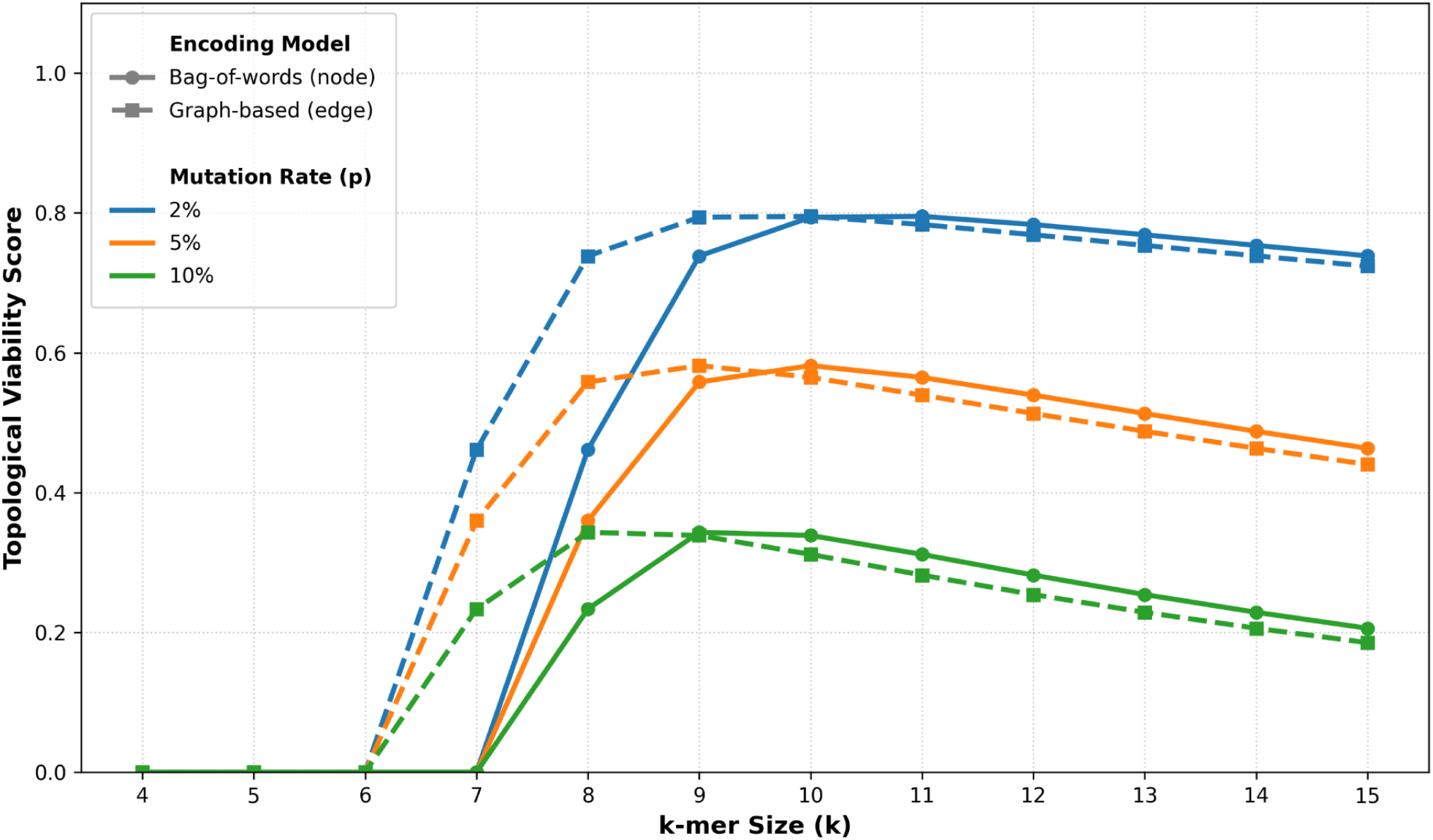
Theoretical Topological Viability of viral sequence graphs under varying evolutionary mutation rates. The Topological Viability, a metric balancing the combinatorial uniqueness of a sequence feature against its probability of surviving evolutionary mutation, is plotted across varying k-mer sizes for a baseline genome length (L=30kb). The models are evaluated at mutation rates (p) corresponding to intra-species variants (2%), species boundaries (5%), and deep evolutionary divergence (10%). The standard bag-of-words model (solid lines) evaluates isolated sequence nodes possessing a biological footprint of k nucleotides. Conversely, the graph-based model employed by HyperSketch (dashed lines) evaluates structural *de Bruijn* transitions with an expanded biological footprint of k+1nucleotides. By plotting both encoding paradigms in a unified space, it demonstrates that the structurally larger footprint of a graph edge causes it to decay faster than an isolated node under equivalent mutation rates. Consequently, to maintain an identical signal-to-noise ratio and maximize the surviving homologous signal, the graph-based sketch fundamentally necessitates an operational parameter exactly one k-mer size lower than the bag-of-words model across all modeled divergence thresholds.

To maintain an identical signal-to-noise ratio, the graph model must compensate for this inherent fragility by uniformly shifting its operational k-mer size to the left. For example, at a standard 5% viral mutation rate, the bag-of-words model maximizes viability at k=10. Conversely, the graph model reaches its absolute apex at k=9. At a slower 2% mutation rate, where signal degradation is less severe, the viability of the graph model plateaus equally across k=9 and k=10. We analytically concluded that k=9 represents the most conservative, broadly applicable optimum for extracting structural topologies across the diverse evolutionary distances inherent to viral phylogenomics.

### Empirical Validation of k=9 as the Topological Optimum

While theoretical models operate under the assumption of uniform nucleotide distribution, real viral genomes contain complex repetitive motifs and GC-biases that alter the true threshold of homoplasy. To definitively validate the theoretical -1 shift in k-mer optimization, we evaluated the absolute biological ground truth by performing exact set intersections across our dataset of ∼26 thousand viral genomes.

To comprehensively profile the sequence features, we evaluated three distinct metrics across varying k-mer footprints:

● <u>Average Unique K-mers:</u> the mean number of distinct k-mers extracted per genome. This quantifies the raw vocabulary size of the sequence.
● <u>Average Common Features (ACF):</u> the absolute mean number of identical k-mers shared between pairwise genome comparisons across the dataset. This is the most critical metric, as it represents the true surviving homologous signal connecting related viruses.
● <u>Information Ratio:</u> the ratio of unique k-mers to the total genome length. This metric represents the specificity of the extracted features. A low ratio indicates rampant sequence repetition (high homoplasy), while a ratio approaching 100% indicates that nearly every extracted k-mer is unique.

Table 1 highlights the critical danger of blindly prioritizing sequence uniqueness when constructing structural sketches. Traditional MinHash estimators, such as Mash, typically favor larger k-mer sizes (e.g., k=12 or higher) to push the Information Ratio past 0.90, ensuring that nearly all extracted features are globally unique. However, our dataset-wide intersections prove that as the footprint expands beyond 10 bases, the ACF, which represents the actual homologous biological signal surviving between related viruses, begins to steadily decay. Using k=12 specifically sacrifices nearly 500 homologous structural features per genome comparison relative to the optimal peak.

**Table 1:** Empirical analysis of sequence features across ∼26 thousand viral genomes. The Information Ratio models uniqueness, while the Average Common Features (ACF) directly measures the surviving homologous signal between genome pairs. The peak ACF across the dataset occurs at a biological footprint of exactly 10 bases.

| k-mer | Average Unique K-mers | Average Common Features | Information Ratio |
| --- | --- | --- | --- |
| 7 | 5,419 | 3,668 | 7.87% |
| 8 | 14,084 | 6,474 | 20.45% |
| 9 | 29,213 | 8,436 | 42.41% |
| 10 | 45,803 | 8,742 | 66.49% |
| 11 | 56,903 | 8,485 | 82.61% |
| 12 | 62,243 | 8,302 | 90.36% |

The empirical ACF peaks flawlessly at a biological sequence footprint of exactly 10 bases (8,742 common features). To operate at this exact biological maximum, a node-based MinHash tool evaluates 10-mers (k=10). However, because HyperSketch maps topological edges, k=9 perfectly encapsulates this 10-base footprint. This empirical intersection data decisively proves that utilizing k=12 for graph-based sketches results in severe topological shattering, and formally validated k=9 as the absolute optimum for structurally mapping viral genomes.

### Hyperparameter Space Sweep and Genomic Distance Estimation

To ensure a rigorous comparison between the structural graphs of HyperSketch and traditional bag-of-words estimators, we established a footprint-normalized baseline against two distinct tools: Mash and HyperGen. Benchmarking against HyperGen serves as a critical ablation study. Because both tools utilize identical underlying HDC algebraic operations, any divergence in their output is strictly attributable to HyperSketch’s novel sequence graph encoding. As established in the previous section, both Mash and HyperGen evaluate isolated nodes and are therefore parametrized at k=10 to exactly match the 10-nucleotide biological footprint of a HyperSketch edge at k=9. This footprint normalization mathematically isolates the core algorithmic data structure as the sole independent variable.

To optimize the remaining HyperSketch variables, and to prove the stability of HDC in genomic applications, we evaluated the dataset across a full combinatorial grid of hyperdimensional states (Supplementary Figure S1). To highlight the impact of the core variables, Figure 2 isolates the absolute parameter extremes against the optimal state in a focused 2x2 grid. We varied the Hypervector Dimensionality (D) across {512, 8192} and the Memory Bank size (M) across {1,8192}. To maintain a strictly fair resource comparison, the Mash sketch size and the HyperGen vectors dimensionality were constrained to perfectly equal the selected HyperSketch dimensionality (D). Furthermore, to facilitate visual clarity and prevent severe overlapping in the density maps, the data points rendered in Figure 2 were randomly downsampled to 100 thousand pairs from the original ∼338 million all-*vs*-all dataset comparisons.

**Figure 2:**
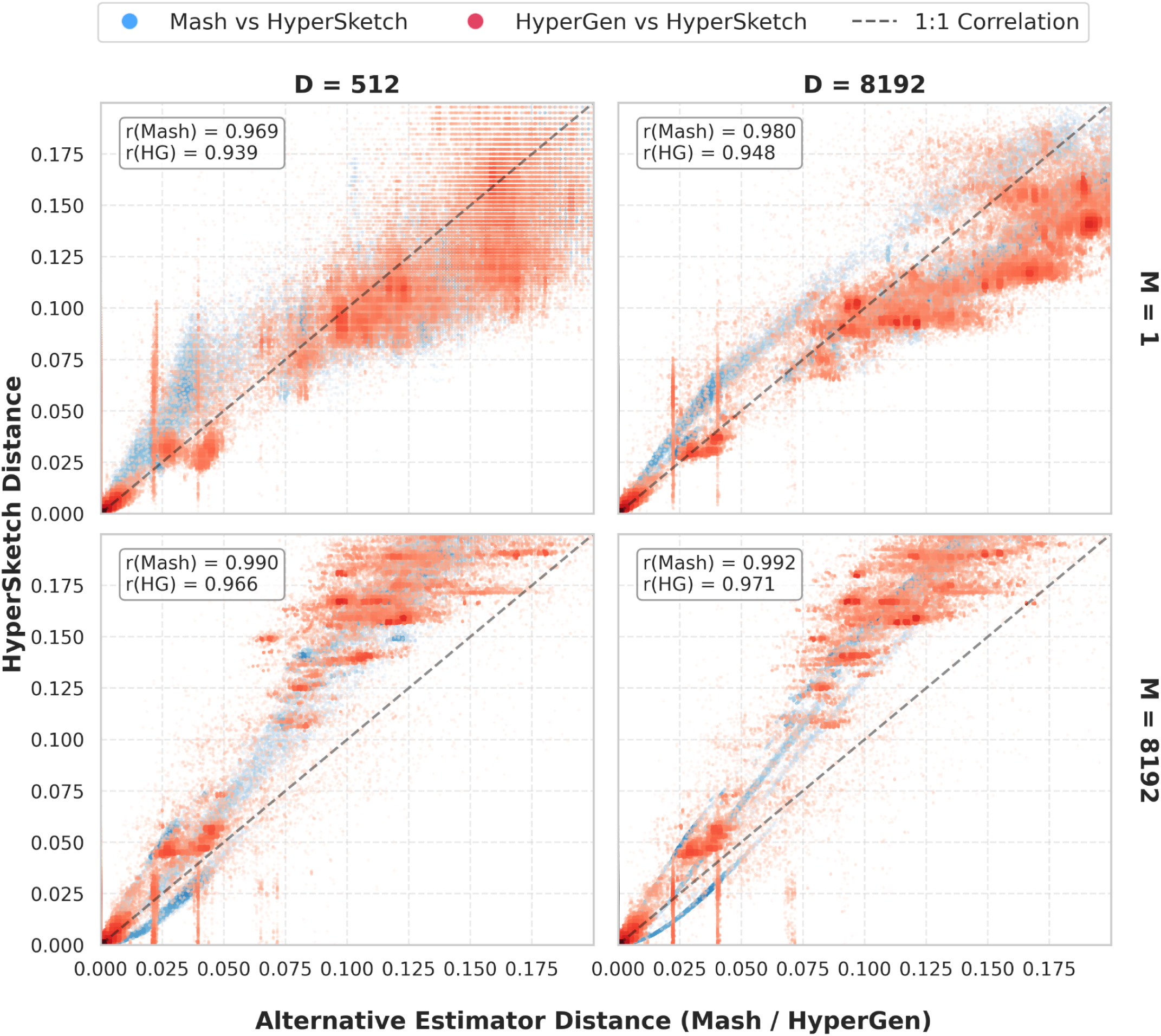
Combinatorial sweep analysis of HyperSketch hyperparameter states against equivalent Mash and HyperGen distance estimations. The columns represent increasing Memory Bank capacity (M), while the rows represent increasing Hypervector Dimensionality (D). Each subplot features a two-color dotplot comparing HyperSketch pairwise distances against Mash (red) and HyperGen (blue). The correlation tightens as hash collisions are mitigated (increasing M) and statistical variance is reduced (increasing D).

Analysis of the combinatorial grid (Figure 2) reveals two distinct axes of architectural stability within the HyperSketch algorithm:

1. <u>The impact of Memory Bank capacity (M):</u> contrasting the top row (M=1) with the bottom row (M=8192) visualizes the effect of hash collisions on the implicit *de Bruijn* graph. When the memory bank is heavily constrained to a single bucket (M=1), every structural edge from the genome collides, resulting in severe destructive interference during the bundling phase. This manifests as a highly dispersed, high-variance similarity cloud. As M increases to 8,192, the HDC associative memory approaches true sparsity, perfectly resolving the signal and pushing the correlations significantly tighter to the diagonal.
2. <u>The impact of Hypervector Dimensionality (D):</u> contrasting the left column (D=512) with the right column (D=8192) validates the fundamental premise of high-dimensional geometry. At lower dimensionalities, the angles between pseudo-orthogonal hypervectors lack resolution, resulting in discrete vertical banding or quantization artifacts during the cosine calculation. As dimensionality increases to 8,192, the geometric resolution smooths into a continuous, high-fidelity gradient.

At the optimal parameter state (k=9, D=8192, and M=8192), HyperSketch correlates strongly with Mash (Pearson correlation r=99.2%) and HyperGen (Pearson correlation r=97.1%). However, as observed in the full supplementary sweep, intermediate constrained memory states (e.g., M=512) actually yield a slightly higher correlation with Mash (r=99.8%). This establishes that compressed, binarized HDC vector bundling can successfully reconstruct whole-genome evolutionary distances on par with established MinHash protocols, while laying the mathematical groundwork to evaluate structural divergence.

Beyond establishing optimal grid parameters, the comparative evaluation in Figure 2 highlights a fundamental mathematical limitation of set-based estimators when evaluating highly divergent genomes. In the standard MinHash framework (utilized by both Mash, HyperGen, and the final distance calculation of HyperSketch), the Jaccard index of intersecting k-mers is converted to a genomic distance using a logarithmic transformation that models a Poisson distribution of point mutations. However, when two genomes are highly divergent and their shared k-mer vocabulary approaches zero, this logarithmic formula yields highly unstable and mathematically unintuitive distance estimates. Previous studies have well-documented this limitation, demonstrating that MinHash-derived ANI estimations become unreliable for distantly related genomes, specifically degrading as ANI falls below 80-85% [31].

Because HyperSketch translates its continuous geometric cosine similarity back into an equivalent Jaccard index to apply this standard mutation model, it mathematically inherits this limitation. At extreme divergences, the geometric similarity of the sketches represents statistical noise (the expected orthogonality of random hypervectors) rather than true biological homology. When stretched by the logarithmic distance formula, this noise inflates into a diffuse band of highly variable distances.

To prevent this instability from introducing analytical artifacts, we applied a strict distance filter across all estimators, explicitly bounding the pairwise comparisons in our analysis to a maximum distance of 0.2 (>=80% ANI). This constraint ensures that the distance calculations remain within the mathematically stable regime of the logarithmic transformation. Consequently, the distinct deviations observed between HyperSketch and the baseline estimators in Figure 2 are strictly attributed to true structural rearrangements and topological variation, rather than algorithmic breakdown at high divergence.

### Graph Reconstruction Fidelity and Signal Degradation

To mathematically explain the correlation boundaries observed in the hyperparameter sweep, we evaluated the internal capacity limits of the HDC Memory Bank. In Vector Symbolic Architectures, a composite memory vector possesses a strict capacity limit. If too many hypervectors are superimposed, destructive interference overwhelms the underlying signals, rendering the original data unrecoverable.

To quantify this, we performed a reconstruction analysis across the 2x2 parameter grid. For a subset of genomes, we extracted the true biological *de Bruijn* edges (source → target) constituting the ground-truth sequence graph. To query the encoded HyperSketch sketch, the source sequence was hashed to identify its spatial row, and the Hamming distance was computed against the deterministically regenerated target hypervector. Because the expected signal strength of a superimposed hypervector degrades proportionally to 1/√*N* (where N represents the collision density of edges mapped to a single bucket), we established a dynamic, bucket-specific detection threshold. To guarantee that a detected match was not a random geometric artifact of high-dimensional space, this dynamic threshold was strictly capped by an absolute 5-sigma statistical noise floor (p<3×10^−7^). Edges whose Hamming distance successfully fell below this threshold were mathematically confirmed as isolated from the background noise. The global reconstruction rate (recall) was then calculated as the percentage of total unique ground-truth edges successfully recovered from the compressed memory matrix.

As demonstrated in Figure 3, the accuracy of HyperSketch’s evolutionary distance estimation is strictly bound by its graph reconstruction fidelity. At highly constrained states (e.g. D=512, M=1), severe hash collisions in the spatial addressing layer cause the reconstruction rate to fall to 0%, meaning the entirety of the structural graph is lost to noise. This signal degradation directly drives the underestimation of genomic distance and the lower baseline correlation (r=0.969) observed in the highly constrained panels of Figure 2. Conversely, as M and D scale towards 8,192, the associative memory approaches sufficient sparsity. The reconstruction rate converges to 100%, perfectly preserving the genome’s structural syntax and allowing the binarized cosine calculation to accurately mirror the true biological distance (r=0.992).

**Figure 3:**
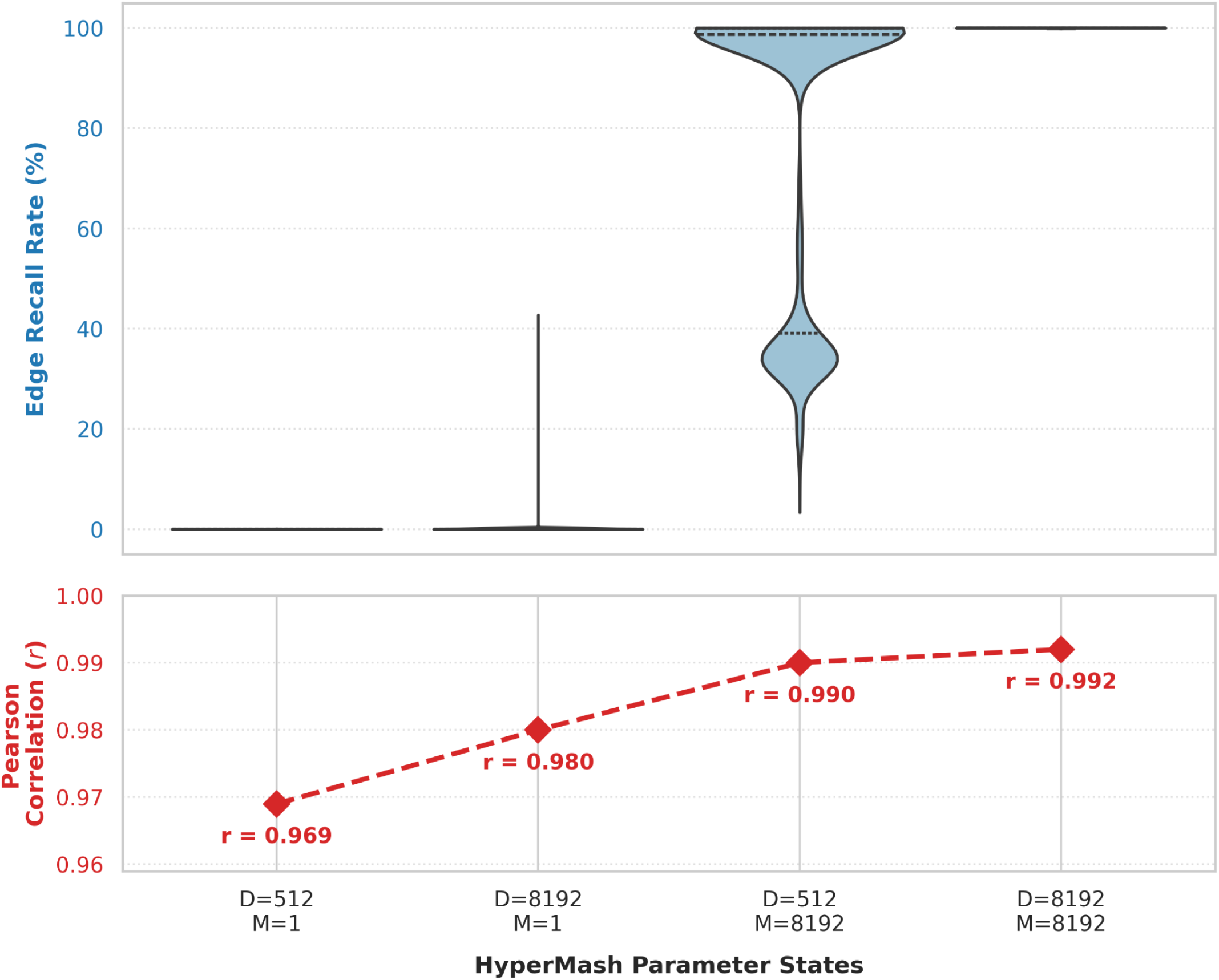
Vertically stacked subplots correlating the HDC graph reconstruction rate with the accuracy of genomic distance estimation across four key parameter states. The top panel (blue violins) represents the probability density of the structural edge recall rate across the evaluated genomes. The bottom panel (red dashed line) tracks the corresponding Pearson correlation (r) of the HyperSketch distance estimates against the baseline compositional metric. As the Memory Bank size (M) and dimensionality (D) increase, destructive interference is mitigated, allowing the true graph topology to be fully recovered and maximizing distance estimation accuracy

### Computational Efficiency and Sketch Footprint

A primary objective of alignment-free sketching is to enable the rapid, memory-efficient clustering of massive, dataset-scale databases. We benchmarked the computational footprint of HyperSketch against Mash and HyperGen to ensure the incorporation of structural graph topology did not sacrifice performance. Benchmarks were conducted utilizing a single thread on a high-performance computing infrastructure with 4 Intel Xeon Platinum 8276 L Central Processing units (CPUs) (112 cores/224 threads) @ 2.20GHz and 6TB of Random Access Memory (RAM) running CentOS Linux 7.

To normalize the operational overhead of disk I/O and isolate the pure algorithmic cost of distance estimation, we executed a controlled batch of 100 viral genomes, resulting in 4,950 pairwise comparisons. The cumulative time was averaged to determine the precise cost of a single pairwise comparison, with the complete operational footprint summarized in Table 2.

**Table 2:**
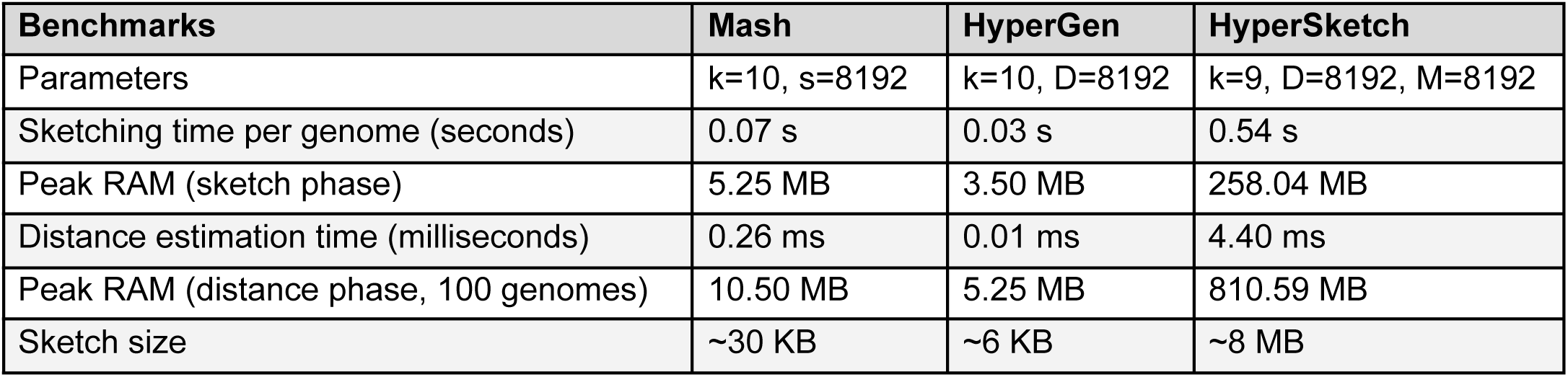
Computational performance benchmarks isolating the resources required to sketch a single viral genome and compute a single pairwise distance. Distance metrics were computed over a 100-genome sub-matrix (4,950 comparisons) on a single CPU thread to effectively average out systemic file-loading latency.

| Benchmarks | Mash | HyperGen | HyperSketch |
| --- | --- | --- | --- |
| Parameters | k=10, s=8192 | k=10, D=8192 | k=9, D=8192, M=8192 |
| Sketching time per genome (seconds) | 0.07 s | 0.03 s | 0.54 s |
| Peak RAM (sketch phase) | 5.25 MB | 3.50 MB | 258.04 MB |
| Distance estimation time (milliseconds) | 0.26 ms | 0.01 ms | 4.40 ms |
| Peak RAM (distance phase, 100 genomes) | 10.50 MB | 5.25 MB | 810.59 MB |
| Sketch size | ~30 KB | ~6 KB | ~8 MB |

#### Single-Genome Encoding Profile

During the initial sketch generation phase, standard bag-of-words estimators (Mash and HyperGen) rapidly process isolated nodes. HyperSketch, however, incurs additional computational overhead due to the structural complexities of graph encoding. Specifically, the framework must extract adjacent *de Bruijn* edges, deterministically generate high-dimensional pseudo-orthogonal vectors for the target nodes, and superimpose them into the spatial matrix via holographic bundling. Consequently, generating a single HyperSketch sketch requires 0.54 seconds, demanding a higher initial encoding cost than the baseline MinHash architecture (0.07 seconds). The peak RAM during this phase (258.04 MB) reflects the exact uncompressed memory allocation (M x D x 4 bytes) required to safely superimpose the topological edges before final quantization.

#### Pairwise Distance Estimation

The computational overhead of graph encoding is decisively recouped during the distance estimation phase. Both HyperSketch and HyperGen leverage identical HDC mechanics, computing geometric similarity via bitwise XOR and popcount instructions. Evaluating a single pair of massive 2D HyperSketch sketches requires only 4.40 milliseconds on a single thread. This ultra-fast binary comparison allows HyperSketch to process structural homologies in fractions of a second.

During the distance phase for 100 genomes, HyperSketch utilized 810.59 MB of RAM, predictably scaling to the ∼218 GB required to hold the entire 26 thousand genome dataset in-memory. Because pairwise distance computations scale seamlessly across multiple processing cores, this lightweight 4.40 ms per-pair baseline permitted our 224-thread infrastructure to resolve the entire ∼338-million comparison matrix in a matter of minutes.

#### Disk Space Complexity and the Topological Trade-off

Capturing the syntax and structural topology of a genome inherently requires a larger memory footprint than storing an unstructured bag-of-words. A standard Mash sketch parametrized to store 8,192 features requires approximately 30 KB per genome, as it relies on a simple 1D array of 64-bit integers. While an exact 10-mer presence bitset would theoretically require only ∼64 KB, such structures strictly represent isolated vocabularies. HyperSketch’s associative memory of size M x D represents a necessary computational trade-off; it is specifically scaled to holographically integrate structural syntax rather than keeping facts separate. For our optimized parameters (D=8192, M=8192), the resulting sketch demands a significantly larger storage footprint of ∼8 MB per genome.

This increase in space complexity represents a fundamental computational trade-off for gaining structural variation awareness. However, to ensure these graph sketches remain practical for large-scale analyses, HyperSketch employs aggressive data quantization. The raw encoding phase utilizes 32-bit integers to accurately superimpose topological edges, which would theoretically require over 268 MB per genome to store on disk. By applying a binarization protocol and densely bit-packing the resulting matrix (storing 8 dimensions per byte), HyperSketch achieves a strict 32-fold compression ratio. This optimization restricts the final associative memory graph to a highly manageable ∼8 MB.

### Distance Scaling and Strain-Level Resolution

While baseline evolutionary distance is largely driven by the steady accumulation of point mutations, viral evolution is heavily punctuated by major architectural events such as homologous recombination, large-scale inversions, and horizontal gene transfer. Because these events shuffle existing genomic content without introducing novel sequences, they present a critical blind spot for purely compositional estimators. We evaluated HyperSketch’s ability to act as a topological sensor to capture this hidden dimension of viral evolution.

To prove the practical utility of this topological sensitivity at the sub-lineage level, we isolated a dataset of 239 *Human mastadenovirus D* (HAdV-D) genomes. HAdV-D is characterized by exceptionally high rates of homologous recombination, frequently swapping major capsid genes (hexon, penton, and fiber) to evade host immune responses [32]. Because these strains are simply exchanging existing viral gene cassettes, their overall k-mer vocabularies remain virtually identical.

We compute the all-vs-all pairwise distance matrices for the HAdV-D dataset using both Mash (k=10, s=8192) and HyperSketch (k=9, D=8192, M=8192) to evaluate their capacity for strain-level resolution. While HyperGen was included in our broader macro-evolutionary benchmarks, it was explicitly excluded from this sub-lineage analysis. Because HyperGen mathematically mirrors Mash by modeling genomes as unstructured bags of k-mers, the two tools yield high congruent, composition-driven distance estimates at micro-evolutionary scales. We therefore utilized Mash as the singular, gold-standard representative for all purely compositional estimators.

As demonstrated in Figure 4A, the purely compositional Mash estimator compresses the evolutionary relationships of the HAdV-D strains into narrow, tightly constrained peaks. Because Mash evaluates the genome as an unstructured bag-of-words, it registers the shared recombinant gene cassettes as nearly identical, minimizing the reported distance.

**Figure 4:**
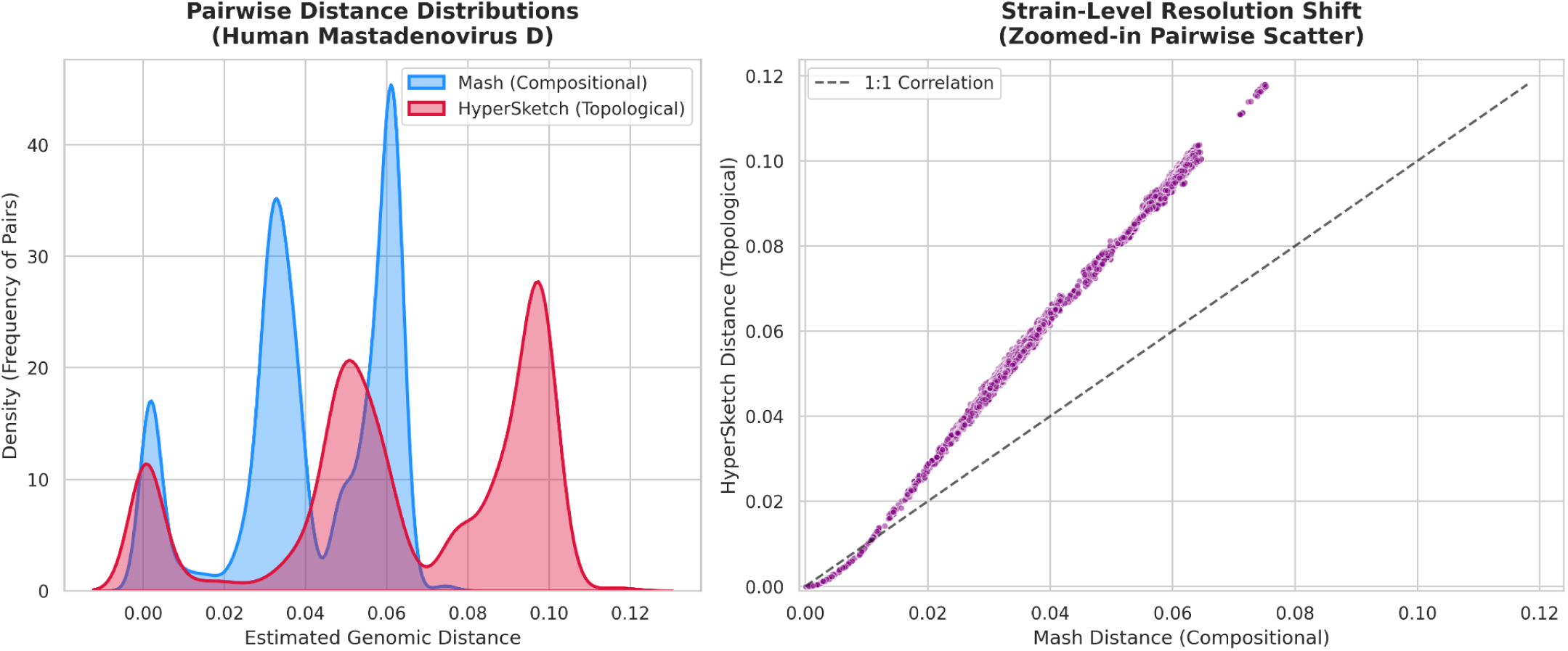
Strain-level resolution and topological magnification in *Human mastadenovirus D*. <u>Panel A</u> – Kernel density estimation of pairwise distances reveals that HyperSketch (red) significantly expands the dynamic range of evolutionary distance estimations compared to the purely compositional Mash baseline (blue). <u>Panel B</u> – A zoomed-in pairwise scatter plot demonstrates that HyperSketch systematically applies a heavier distance penalty to closely related strains, reflecting the added geometric cost of severed topological edges resulting from homologous recombination.

On the other hand, the distance formulation of HyperSketch naturally introduces a scaling effect. Because the transformation scales with the expanded k+1 footprint, it mathematically expands the dynamic range of the distance metric, stretching the density distribution into wider, more highly resolved sub-lineage peaks (Figure 4A). This scaling is explicitly visualized in the pairwise scatter plot (Figure 4B). At the micro-evolutionary scale, every pairwise comparison strictly deviates above the 1:1 correlation baseline. By layering this expanded HyperSketch distance alongside traditional compositional metrics, researchers can achieve higher-resolution sub-typing for closely related viral strains.

### Quantitative Assessment of Strain-Level Resolution

To quantitatively validate the topological magnification observed in *Human mastadenovirus D* across a broader evolutionary context, we evaluated the capacity of both HyperSketch and Mash to resolve known strain-level taxonomies directly from raw genomic distances. Utilizing high-quality metadata from the NCBI Dataset API [33], we isolated viral species exhibiting robust, multi-genome strain annotations. To ensure the clustering metrics were evaluating true biological groupings rather than database artifacts, we strictly filtered the dataset to exclude species where NCBI metadata artificially cataloged every sequenced isolate as a unique strain (enforcing a minimum threshold of 1.5 genomes per true strain).

For each of the strictly curated species, we performed agglomerative hierarchical clustering across a high-resolution sweep of continuous distance thresholds (translating to 90.0% – 100.0% ANI). We measured the topological accuracy of the resulting dendrograms against the NCBI ground truth using the Adjusted Rand Index (ARI) [34]. Furthermore, to assess the geometric clarity of the un-thresholded raw distance space, we computed the Silhouette coefficient [35], which measures how tightly cohesive identical strains are relative to their geometric distance from different strains.

The clustering results (Table 3) demonstrate that embedding the *de Bruijn* graph topology natively into the sketch does not compromise baseline compositional accuracy, while significantly enhancing the cluster separation of diverging strains in specific contexts.

**Table 3:** Quantitative comparison of clustering accuracy (ARI) and structural geometric separation (Silhouette score) between HyperSketch (HM) and Mash across highly-curated viral species.

| Species | Genomes | True Strains | HM Max ARI | Mash Max ARI | HM Silhouette | Mash Silhouette | Delta Silhouette |
| --- | --- | --- | --- | --- | --- | --- | --- |
| *Plum pox virus* | 11 | 7 | 1.000 | 1.000 | 0.302 | 0.261 | 0.041 |
| *Gequatrovirus G4* | 152 | 14 | 0.996 | 0.999 | 0.913 | 0.890 | 0.023 |
| *Vaccinia virus* | 60 | 32 | 0.585 | 0.585 | 0.035 | 0.042 | -0.007 |
| *Papaya leaf curl virus* | 12 | 8 | 0.101 | 0.101 | -0.114 | -0.093 | -0.021 |
| *Affertcholeramvirus CTXphi* | 11 | 3 | 0.481 | 0.481 | 0.236 | 0.289 | -0.053 |

For example, in *Plum pox virus*, both Mash and HyperSketch perfectly reconstructed the true strain taxonomy at their optimal thresholds, achieving an ARI of 1.0. However, evaluating the continuous distance matrices revealed that HyperSketch generated a geometrically superior separation space, achieving a Silhouette score of 0.302 compared to Mash’s 0.261.

This distance scaling effect was similarly pronounced in our largest viable dataset, *Gequatrovirus G4* (152 genomes). While both tools achieved near-perfect strain classification (ARI > 0.99), HyperSketch produced a more highly defined structural distance space with a Silhouette score of 0.913, notably outperforming the purely compositional metric (0.890).

Conversely, for species such as *Vaccinia virus* and *Affertcholeramvirus CTXphi*, the purely compositional Mash estimator achieved slightly higher Silhouette scores despite parity in ARI. This divergence perfectly encapsulates the operational distinction between the two estimators. If strains within a specific viral lineage differ exclusively through isolated point mutations (SNPs) while sharing the exact same structural genome arrangement, pure k-mer composition tracks that localized decay smoothly. However, when structural divergence, such as genomic rearrangements, indels, or distinct graph topologies, defines the evolutionary split, the mathematical scaling of HyperSketch provides enhanced empirical separation.

These metrics confirm that the wider dynamic range observed in HyperSketch distances improves clustering performance. By anchoring k-mer content within a hyperdimensional *de Bruijn* topology and applying a steeper logarithmic transformation, HyperSketch pulls closely related strains further apart in the distance space while maintaining tight cohesion among highly conserved isolates. Consequently, HyperSketch provides a higher-resolution geometric metric for defining novel taxonomic clusters in datasets where optimal similarity thresholds are not known *a priori*.

While the observed topological magnification aligns with the known high rates of homologous recombination in *Human mastadenovirus D*, a limitation of the current study is the absence of independent, alignment-derived breakpoint validation. Further work will necessitate the use of simulated structural variants to rigorously quantify the exact sensitivity of the sketch to specific genomic rearrangements.

## DISCUSSION AND CONCLUSIONS

The exponential growth of genomic data requires comparative methods that are not only computationally efficient but also biologically expressive. While MinHash-based estimators have revolutionized the field by reducing genomes to compact k-mer sets, their bag-of-words approach inherently discards the structural context of the genome. In this study, we introduced HyperSketch, a novel sketching algorithm that bridges this gap by leveraging HDC to encode the topology of a genome’s *de Bruijn* graph into a fixed-size, continuous hypervector space.

By binding k-mers to their adjacent edges prior to bundling, HyperSketch successfully preserves the sequence of elements that purely compositional methods ignore. Our results demonstrate that this structural embedding yields a highly sensitive distance metric. Unlike Mash and HyperGen, HyperSketch utilizes an adjusted logarithmic transformation that scales with the k+1 biological footprint of its structural edges. This mathematical formulation introduces a scaling effect that expands the dynamic range of distance estimates. Consequently, HyperSketch yields distance estimations that magnify the separation between closely related lineages, effectively improving the signal-to-noise ratio of the separation space at micro-evolutionary scales.

Our quantitative assessment across highly-curated viral datasets validated the practical utility of this magnification. While HyperSketch maintained baseline compositional clustering accuracy, achieving parity with Mash in reconstructing known taxonomic relationships (ARI), it significantly improved the geometric clarity of the resulting distance space. In complex, multi-strain datasets such as *Plum pox virus* and *Gequatrovirus G4*, HyperSketch produced substantially higher Silhouette scores than pure k-mer estimators. This proves that HyperSketch pulls structurally distinct strains further apart while maintaining tight geometric cohesion among highly conserved isolates, effectively improving the signal-to-noise ratio of the separation space.

These findings clearly delineate the operational niches for compositional versus topological estimators. If strains within a specific lineage differ exclusively through isolated point mutations while sharing identical structural arrangements, purely compositional metrics like Mash remain highly efficient and accurate. However, in evolutionary contexts where structural divergence defines the taxonomic split, pure k-mer composition fails to capture the full biological reality. In these scenarios, HyperSketch provides a mathematically distinct clustering space. By expanding the dynamic range of distance estimates, HyperSketch provides a higher-resolution metric that is particularly advantageous for defining novel taxonomic clusters, grouping uncharacterized environmental isolates, or establishing boundaries where optimal ANI thresholds are not known *a priori*.

The transition from k-mer sets to graph-based hypervectors introduces a computational tradeoff, requiring larger sketch sizes and increased computational overhead compared to highly optimized MinHash implementations. However, the inherent properties of HDC, specifically, its reliance on highly parallelizable, low-precision vector operations, make HyperSketch an ideal candidate for future hardware acceleration. As processing-in-memory (PIM) and neuromorphic architectures become more prevalent, the bottleneck of high-dimensional bundling will diminish, allowing topological sketching to operate at the speed of current compositional methods [36].

Furthermore, while HyperSketch is currently implemented as a standalone C++ utility to maximize processing speed, future extensions will explore cross-language bindings to interface seamlessly with existing Python-based HDC ecosystems, such as *hdlib* [37,38], facilitating the integration of structurally-aware genomic sketches into broader machine learning workflows.

Future comparative assessments will benefit from benchmarking against alternative order-aware sketches, such as Order Min Hash, and utilizing strictly held-out taxonomic splits to prevent parameter optimization bias. Additionally, incorporating alternative concordance metrics beyond linear correlation will help further dissect the scale shifts observed in structural divergence.

In conclusion, HyperSketch demonstrates that genomic sketching does not have to be limited to structural abstraction. By mapping the *de Bruijn* graph into hyperdimensional space, we provide a mathematically continuous, structurally-aware metric that enhances the resolution of genomic comparisons, paving the way for more nuanced, scalable analyses of structural variation and microbial evolution.

## ADDITIONAL INFORMATION

### Abbreviations

ACF: Average Common Features
ANI: Average Nucleotide Identity
ARI: Adjusted Rand Index
CPU: Central Processing Unit
ENA: European Nucleotide Archive
HDC: Hyperdimensional Computing
MAP: Multiply-Add-Permute
NCBI: National Center for Biotechnology Information
PIM: Processing-In-Memory
PRNG: Pseudo-Random Number Generator
RAM: Random Access Memory
VSA: Vector Symbolic Architectures

### Availability

HyperSketch code is open-source and available on GitHub at https://github.com/cumbof/hypersketch under the MIT license.

### Author Contribution

FC conceived the research, designed the methodology, developed the HyperSketch software, and performed the analysis; KD contributed to the software development, data validation, and data curation; MHN, SA, and DB provided expert guidance on the Hyperdimensional Computing architecture, parameter optimization, and hardware considerations; DB supervised the research; all authors contributed to interpreting the results, drafting the manuscript, and have read, revised, and approved the final version.

## Acknowledgments

The authors would like to acknowledge the use of a Large Language Model (Google’s Gemini 3 Pro) for its assistance in rephrasing sentences and improving the overall clarity of the manuscript’s prose. All scientific ideas, mathematical formulations, the algorithmic framework, and software implementations presented in this work are the exclusive product of the authors’ experience and expertise.

## COMBINATORIAL HYPERPARAMETER SWEEP ANALYSIS

To systematically map the operational boundaries and evaluate the algorithmic stability of the HyperSketch encoding framework, we performed an empirical combinatorial parameter sweep analysis (Supplementary Figure S1). Pairwise distance estimations were computed across 30 distinct hyperparameter configurations, iterating the Hypervector Dimensionality (D) across {512, 1024, 2048, 4096, 8192} and the Memory Bank size (M) across {1, 512, 1024, 2048, 4096, 8192}. For each configuration, the Mash sketch size and the HyperGen vector dimensionality were constrained to equal D.

### Influence of Associative Memory Capacity on Signal Integrity (Rows, M)

The progression of rows in Supplementary Figure S1 illustrates how the scale of the Memory Bank dictates the algorithm’s sensitivity to structural variation.

At the extreme minimum (M=1), the entire sequence graph is forced into a single superposition. This collision threshold produces a diffuse distribution where many individual topological signals are lost to destructive interference.

As the array expands to its maximum evaluated capacity (M=8192), the mapping approaches sufficient sparsity to preserve the precise order of encoded topological edges.

### Dimensionality and Geometric Resolution (Columns, D)

The progression of columns demonstrates the relationship between hypervector dimensionality and the continuity of the resulting distance metric.

At lower dimensional constraints (e.g., D=512), the high-dimensional space lacks enough angular capacity to represent fine-grained geometric similarities, producing distinct striations or banding artifacts, introducing localized estimation errors.

Expanding the hypervector dimensionality directly increases the available angular resolution. As D scales toward 8,192, these discrete quantization artifacts are systematically smoothed, resolving into a continuous geometric gradient that enables precise distance estimation across the entire evaluated evolutionary spectrum.

---

**Supplementary Figure S1:**
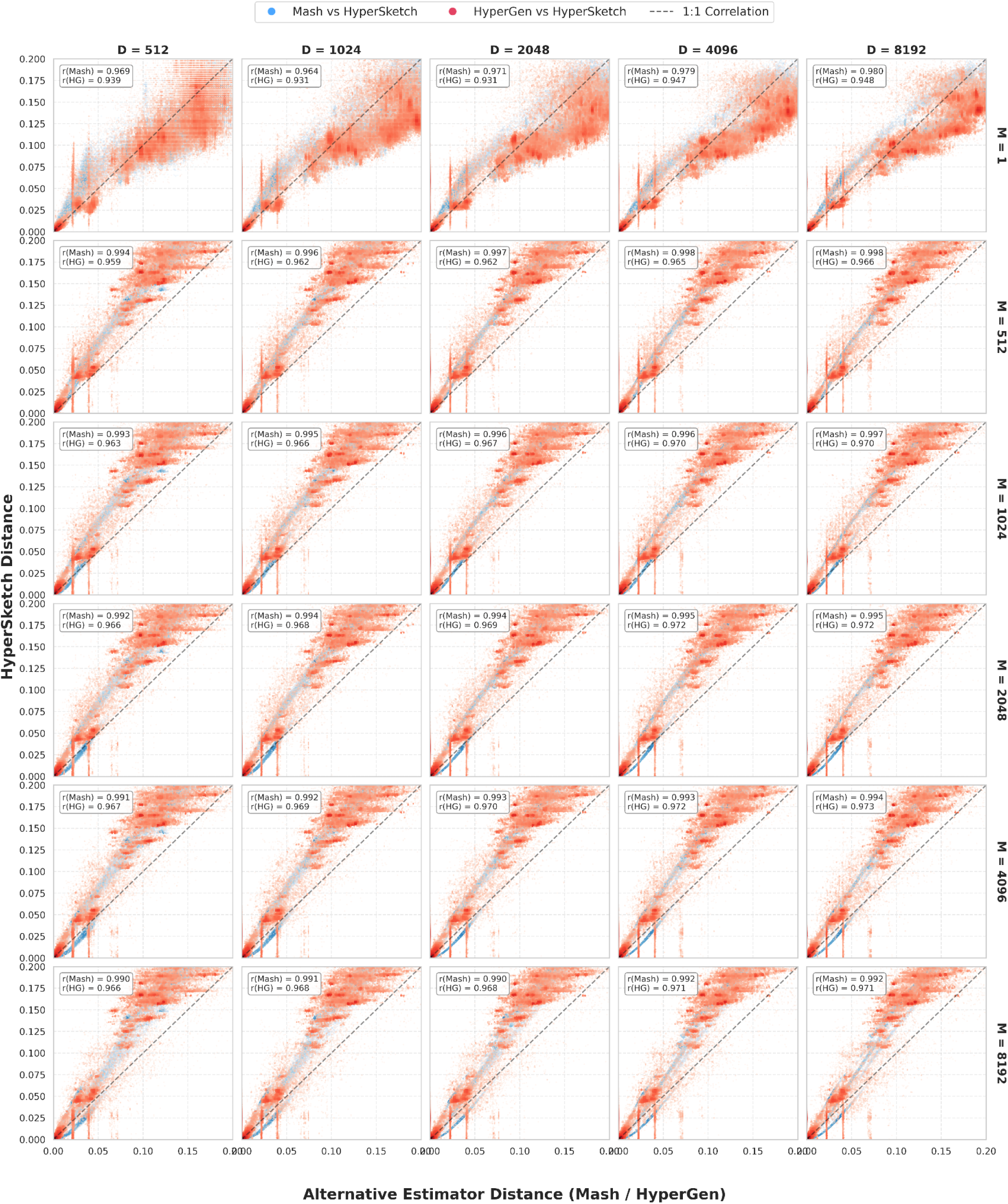
Combinatorial parameter sweep analysis evaluating the effects of Hypervector Dimensionality (D) and Memory Bank size (M) on genomic distance estimations. Scatter plots display a random subsample of 100 thousand pairwise comparisons drawn from the full viral dataset. The Y-axis represents the HyperSketch distance, plotted against the corresponding Mash (blue) and HyperGen (red) distances on the X-axis. The data illustrates the progressive mitigation of discrete quantization artifacts across increasing columns (D), and the transition from topological interference to high-fidelity structural resolution across increasing rows (M).

## REFERENCES

1. Park ST, Kim J. Trends in Next-Generation Sequencing and a New Era for Whole Genome Sequencing. Int Neurourol J. 2016;20: S76–83. doi:10.5213/inj.1632742.371

2. Chaudhari JK, Pant S, Jha R, Pathak RK, Singh DB. Biological big-data sources, problems of storage, computational issues, and applications: a comprehensive review. Knowledge and Information Systems. 2024;66: 3159–3209. doi:10.1007/s10115-023-02049-4

3. Li Y, Chen L. Big Biological Data: Challenges and Opportunities. Genomics, Proteomics & Bioinformatics. 2014;12: 187–189. doi:10.1016/j.gpb.2014.10.001

4. Altschul SF, Gish W, Miller W, Myers EW, Lipman DJ. Basic local alignment search tool. Journal of Molecular Biology. 1990;215: 403–410. doi:10.1016/s0022-2836(05)80360-2

5. Delcher AL, Salzberg SL, Phillippy AM. Using MUMmer to identify similar regions in large sequence sets. Curr Protoc Bioinformatics. 2003;Chapter 10: Unit 10.3. doi:10.1002/0471250953.bi1003s00

6. Saada B, Zhang T, Siga E, Zhang J, Magalhães Muniz MM. Whole-Genome Alignment: Methods, Challenges, and Future Directions. Applied Sciences. 2024;14: 4837. doi:10.3390/app14114837

7. Zielezinski A, Vinga S, Almeida J, Karlowski WM. Alignment-free sequence comparison: benefits, applications, and tools. Genome Biol. 2017;18: 186. doi:10.1186/s13059-017-1319-7

8. Vinga S, Almeida J. Alignment-free sequence comparison—a review. Bioinformatics. 2003;19: 513–523. doi:10.1093/bioinformatics/btg005

9. Zielezinski A, Girgis HZ, Bernard G, Leimeister C-A, Tang K, Dencker T, et al. Benchmarking of alignment-free sequence comparison methods. Genome Biol. 2019;20: 144. doi:10.1186/s13059-019-1755-7

10. Ferraro Petrillo U, Sorella M, Cattaneo G, Giancarlo R, Rombo SE. Analyzing big datasets of genomic sequences: fast and scalable collection of k-mer statistics. BMC Bioinformatics. 2019;20: 138. doi:10.1186/s12859-019-2694-8

11. Das A, Schatz MC. Sketching and sampling approaches for fast and accurate long read classification. BMC Bioinformatics. 2022;23: 452. doi:10.1186/s12859-022-05014-0

12. Sánchez-Reyes A, Fernández-López MG. Sketched reference databases for genome-based taxonomy and comparative genomics. Braz J Biol. 2022;84: e256673. doi:10.1590/1519-6984.256673

13. Ondov BD, Treangen TJ, Melsted P, Mallonee AB, Bergman NH, Koren S, et al. Mash: fast genome and metagenome distance estimation using MinHash. Genome Biol. 2016;17: 132. doi:10.1186/s13059-016-0997-x

14. Jaccard P. Étude comparative de la distribution florale dans une portion des Alpes et du Jura. 1901. doi:10.5169/SEALS-266450

15. Zhang T, Yin Z, Xu X, Yan L, Zhu F, Duan X, et al. RabbitSketch: a high-performance sketching library for genome analysis. Bioinformatics. 2025;41. doi:10.1093/bioinformatics/btaf249

16. Baker DN, Langmead B. Dashing: fast and accurate genomic distances with HyperLogLog. Genome Biol. 2019;20: 265. doi:10.1186/s13059-019-1875-0

17. Liu S, Koslicki D. CMash: fast, multi-resolution estimation of k-mer-based Jaccard and containment indices. Bioinformatics. 2022;38: i28–i35. doi:10.1093/bioinformatics/btac237

18. Kanerva P. Hyperdimensional computing: An algebra for computing with vectors. Advances in Semiconductor Technologies. Wiley; 2022. pp. 25–42. doi:10.1002/9781119869610.ch2

19. Kanerva P. Hyperdimensional computing: An introduction to computing in distributed representation with high-dimensional random vectors. Cognit Comput. 2009;1: 139–159. doi:10.1007/s12559-009-9009-8

20. Cumbo F, Chicco D. Hyperdimensional computing in biomedical sciences: a brief review. PeerJ Comput Sci. 2025;11: e2885. doi:10.7717/peerj-cs.2885

21. Cumbo F, Chicco D, Aygun S, Blankenberg D. Designing vector-symbolic architectures for biomedical applications: ten tips and common pitfalls. PeerJ Comput Sci. 2026;12: e3682. doi:10.7717/peerj-cs.3682

22. Xu W, Hsu P-K, Moshiri N, Yu S, Rosing T. HyperGen: Compact and Efficient Genome Sketching using Hyperdimensional Vectors. Bioinformatics. 2024;40. doi:10.1093/bioinformatics/btae452

23. Poduval P, Alimohamadi H, Zakeri A, Imani F, Najafi MH, Givargis T, et al. GrapHD: Graph-Based Hyperdimensional Memorization for Brain-Like Cognitive Learning. Front Neurosci. 2022;16: 757125. doi:10.3389/fnins.2022.757125

24. Cumbo F, Dhillon K, Joshi J, Chicco D, Aygun S, Blankenberg D. A novel Vector-Symbolic Architecture for graph encoding and its application to viral pangenome-based species classification. BioData Min. 2026. doi:10.1186/s13040-026-00561-1

25. Widynski B. Middle-Square Weyl Sequence RNG. arxiv. 2017. Available: http://arxiv.org/abs/1704.00358

26. Plate TA. Holographic reduced representations. IEEE Trans Neural Netw. 1995;6: 623–641. doi:10.1109/72.377968

27. Pisier G. Grothendieck’s theorem, past and present. Bull New Ser Am Math Soc. 2012;49: 237–323. doi:10.1090/s0273-0979-2011-01348-9

28. Ochiai A. Zoogeographical Studies on the Soleoid Fishes found in Japan and its Neighbouring Regions-I. Nippon Suisan Gakkai Shi. 1957;22: 522–525. doi:10.2331/suisan.22.522

29. Pornputtapong N, Acheampong DA, Patumcharoenpol P, Jenjaroenpun P, Wongsurawat T, Jun S-R, et al. KITSUNE: A Tool for Identifying Empirically Optimal K-mer Length for Alignment-Free Phylogenomic Analysis. Front Bioeng Biotechnol. 2020;8: 556413. doi:10.3389/fbioe.2020.556413

30. Sayers EW, Beck J, Bolton EE, Brister JR, Chan J, Connor R, et al. Database resources of the National Center for Biotechnology Information in 2025. Nucleic Acids Research. 2024;53: D20–D29. doi:10.1093/nar/gkae979

31. Jain C, Rodriguez-R LM, Phillippy AM, Konstantinidis KT, Aluru S. High throughput ANI analysis of 90K prokaryotic genomes reveals clear species boundaries. Nat Commun. 2018;9: 5114. doi:10.1038/s41467-018-07641-9

32. Robinson CM, Singh G, Lee JY, Dehghan S, Rajaiya J, Liu EB, et al. Molecular evolution of human adenoviruses. Sci Rep. 2013;3: 1812. doi:10.1038/srep01812

33. O’Leary NA, Cox E, Holmes JB, Anderson WR, Falk R, Hem V, et al. Exploring and retrieving sequence and metadata for species across the tree of life with NCBI Datasets. Sci Data. 2024;11: 732. doi:10.1038/s41597-024-03571-y

34. Hubert L, Arabie P. Comparing partitions. Journal of Classification. 1985;2: 193–218. doi:10.1007/BF01908075

35. Rousseeuw PJ. Silhouettes: A graphical aid to the interpretation and validation of cluster analysis. Journal of Computational and Applied Mathematics. 1987;20: 53–65. doi:10.1016/0377-0427(87)90125-7

36. Karunaratne G, Le Gallo M, Cherubini G, Benini L, Rahimi A, Sebastian A. In-memory hyperdimensional computing. Nature Electronics. 2020;3: 327–337. doi:10.1038/s41928-020-0410-3

37. Cumbo F, Weitschek E, Blankenberg D. hdlib: A Python library for designing Vector-Symbolic Architectures. J Open Source Softw. 2023;8: 5704. doi:10.21105/joss.05704

38. Cumbo F, Dhillon K, Blankenberg D. hdlib 2.0: Extending Machine Learning Capabilities of Vector-Symbolic Architectures. arxiv. 2026. Available: http://arxiv.org/abs/2601.02509

